# CellClear: Enhancing Single-cell RNA Data Quality via Biologically-Informed Ambient RNA Correction

**DOI:** 10.1101/2024.08.05.606571

**Authors:** Wanxiang Huang, LongFei Hu, Zhengzong Qian, Jue Fan

## Abstract

Ambient RNA, caused by the pool of mRNA molecules released from lysed or dead cells in samples, poses persistent challenges to single-cell RNA sequencing data analysis, and leads to potential problems such as inaccurate classification of cell types and downstream functional analyses. Here we propose CellClear to estimate and remove the ambient RNA expression programs in single-cell RNA sequencing data. CellClear demonstrates significant increase in the accuracy of ambient gene expression correction comparing with state-of-art methods, without distorting native information of individual cell types.

## Background

Single-cell RNA sequencing (scRNA-seq) is a transformative advancement that unveils cellular heterogeneity within complex tissues at single-cell resolution. It facilitates the identification of rare cell types, and allows for the characterization of dynamic cellular states [1,2]. Despite its potentials, scRNA-seq technology can suffer from technical noises, such as low RNA capture efficiency and the presence of ambient RNA [3,4]. Ambient RNA originating from ruptured, dead, or dying cells, as well as cell-free RNA including exome RNA, results in non-zero counts in cell-free barcodes and non-specific gene expressions in cell-associated barcodes [5].

Currently, various tools have been developed for ambient RNA correction. DecontX assumes cells as mixtures of native and contamination-derived counts and iteratively refines their relative proportions [4]. SoupX identifies cluster-specific markers to estimate the fraction of Unique Molecular Identifiers (UMIs) originating from the background and use it to represent the ambient RNA fraction. It is recommended to complement this automated estimation with a set of contamination-related markers for manual estimation [6]. CellBender, a deep generative model based on a neural network, performs well in most datasets [5], but demands significantly high computational costs [7,8]. However, with the increasing trend of analyzing large scale scRNA-seq datasets, there is an urgent need for a method that can not only rapidly and efficiently assess the necessity for decontamination during the quality control stage without prior knowledge, but also accurately correct ambient expression across various data situations such as complex contamination sources.

Here, we introduce CellClear, which can accurately identify and correct ambient genes while preserving the biological features of the data. The corrected data shows a significant improvement of sensitivity during the downstream function analyses. To avoid unnecessary corrections, CellClear also provides an ambient expression level as a QC metric to guide researchers in deciding whether to apply the correction. Comparing to existing methods, CellClear shows superior performance in ambient RNA correction, making it an efficient tool to be included in standard scRNA data analysis pipelines.

## Results and Discussion

In order to address the issue of ambient gene expression in data, we developed CellClear, a tool which is capable of accurately identifying and correcting ambient genes. The performance of CellClear was initially validated using a human-mouse mixed sample and twenty-four human skin samples. This was done to assess its ability to capture ambient genes and to improve cell type classification, respectively. Subsequently, three mouse kidney datasets which included genotype-based ambient expression estimates were introduced to be corrected by CellClear and the other three existing tools to demonstrate the superior performance of CellClear in ambient RNA correction. Finally, one human PBMC sample and two human testis samples were used to investigate the impact of ambient correction tools on downstream analyses, thereby highlighting CellClear’s significant advantages in uncovering biological insights and restoring biological features. Furthermore, in order to evaluate the versatility of CellClear across a diverse range of datasets, a total of 35 samples, including those previously mentioned, were employed in a comprehensive assessment of CellClear. This encompassed the presentation of pertinent information such as sample size, analysis duration, ambient expression levels, and ambient genes (see Additional file 1: Table S1).

### Identification of Ambient Genes in Mixed-Species scRNA-seq Datasets

The “species-mixing” experiments from 10x Genomics, which combine mouse and human cell lines, were employed as the initial dataset to demonstrate CellClear’s capacity to eliminate ambient expression in the presence of ground truth. In this dataset, both human and mouse cells exhibit a considerable number of mouse-specific and human-specific gene counts respectively. However, the majority of exogenous transcripts were successfully identified and removed by CellClear, resulting in a distinct separation between the two species in the UMAP reduction plot (Figure 2A). In the absence of prior knowledge, 10 out of the 20 top ambient genes identified by CellClear were confirmed through post-hoc validation, consisting of mouse genes ranked in the top expression in human cells (e.g., mm10-Rps19) and human genes ranked in the top expression in mouse cells (e.g., hg19-RPS2). (Additional file 3: Table S1). This identification enabled CellClear to successfully eliminate 99.7% of mouse gene counts in human cells and 71.6% of human gene counts in mouse cells, thereby restoring a more purified cell population (Figure 2B).

**Figure 1.**
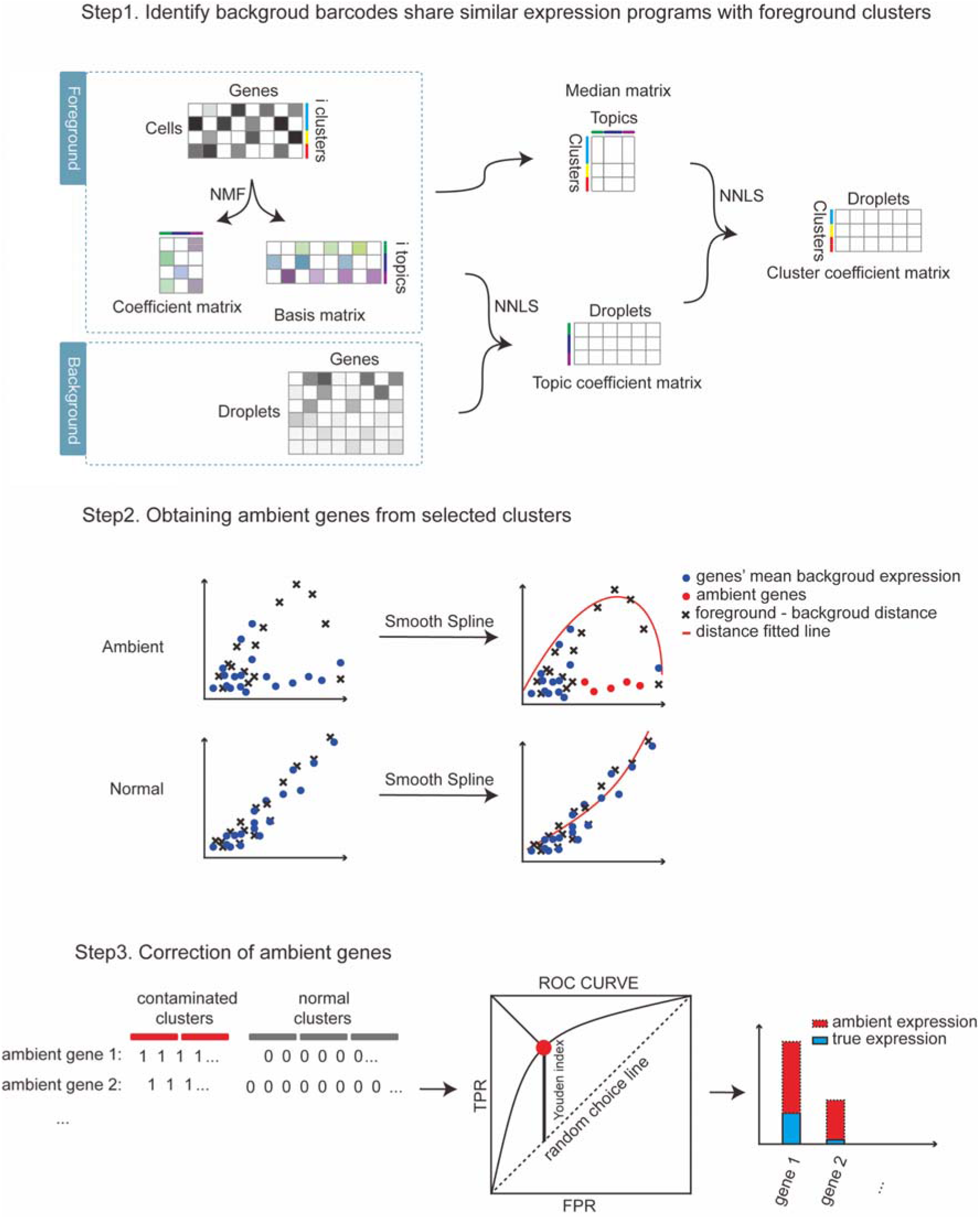
Overview of the CellClear method. Step1. Non-Negative Matrix Factorization (NMF) and two-step Non-Negative Least Squares (NNLS) optimisation are applied to foreground and background matrices. Step2. Genes with abnormal foreground-background distances are taken as ambient genes. Step3. Area Under the Receiver Operating Characteristic Curve (AUC-ROC) and Youden index are used to determine the threshold for each ambient gene.

**Figure 2.**
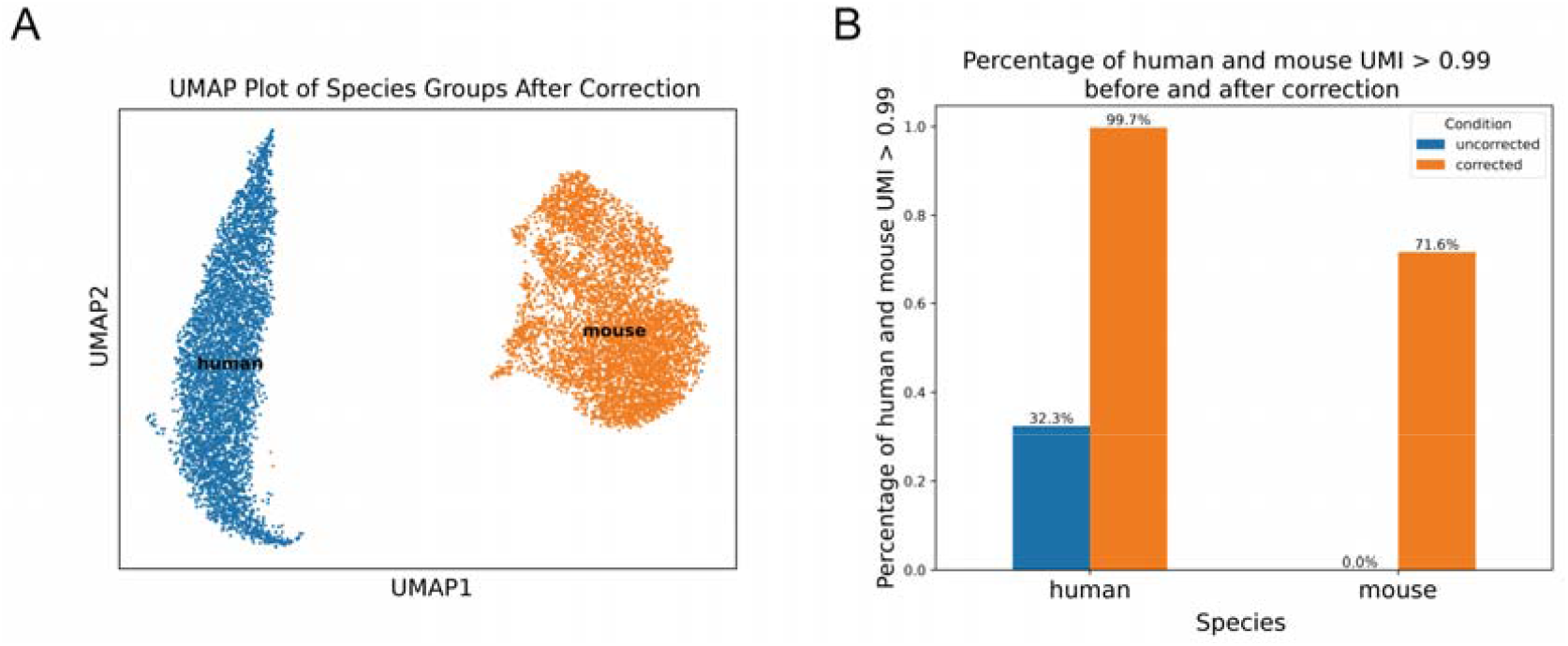
Performance Evaluation of CellClear on the human-mouse cell mixture dataset. (A) UMAP scatter plot shows the destinction of human cells (in blue) and mouse cells (in orange). (B) Percentage of human cells where human gene counts exceed 99%, and the percentage of mouse cells where mouse gene counts exceed 99%, before and after correction.

### Enhancing Cell Type Classification with CellClear in Human Skin Samples

Given the prevalence of ambient expression, which can lead to aberrant cell clustering [9] and inaccurate classification of cell types during the integration of multiple samples, this study sought to evaluate the effectiveness of CellClear in this context. We integrated 24 corrected and 24 uncorrected samples of human skin separately, and annotated them according to the markers provided in the article [10]. In the uncorrected dataset, markers associated with Keratinocytes exhibited significant ambient expression, while markers related to other major cell types demonstrated high specificity for their respective classifications (Figure 3A). This indicates that the high degree of specificity demonstrated by the major cell type markers may serve to mitigate the influence of ambient expression on their annotation.

**Figure 3.**
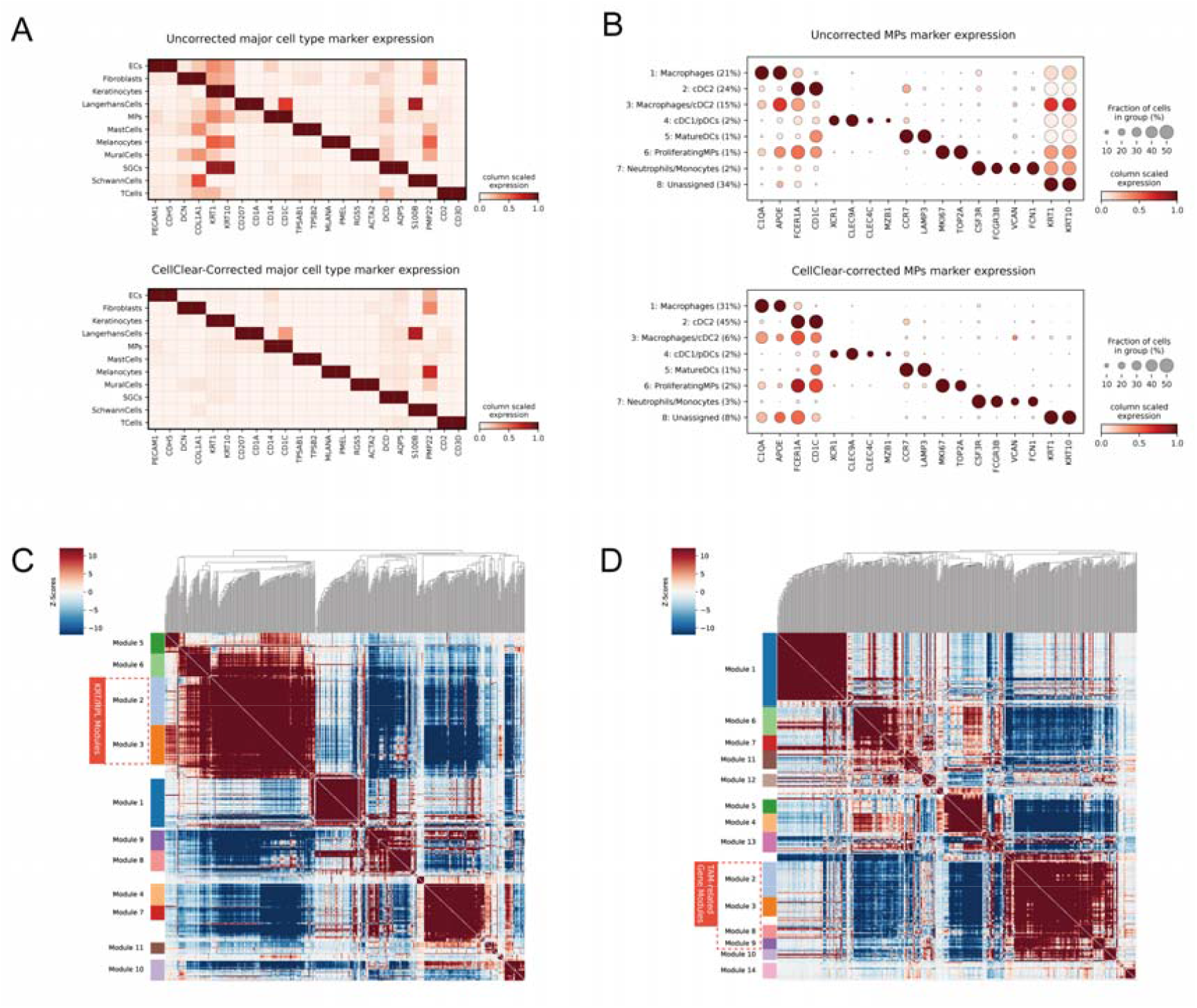
Performance Evaluation of CellClear on the human skin datasets. (A) Matrix Plot shows the expression of markers for each major cell type before and after correction. (B) Dot Plot displays the expression of markers for Mononuclear Phagocytes (MPs) subtypes before and after correction, along with the proportion of each subtype. (C)(D) Heatmap of representative modules of MPs expression before and after correction.

However, when the major cell types were subjected to sub-clustering, it became evident that the pervasive presence of KRT genes exerted a considerable influence on the clustering process. For example, during the sub-clustering of Mononuclear Phagocytes (MPs), approximately 34% of the cells were clustered together (annotated as ‘Unassigned’) without clear identification (Figure 3B). In this instance, the guidance provided by the markers related to MPs, which exhibited lower expression levels, was overshadowed by the markers related to Keratinocytes. Following the application of the CellClear correction, the proportion of ambiguous cells (low expression of classical MPs markers and high expression of KRT genes) was reduced to 8%. Concurrently, the proportions of Macrophages and cDC2 cells markedly increased, rising from 21% and 24% to 31% and 45%, respectively. This further enhanced the classification precision of the subgroups and facilitated a more nuanced understanding of cellular diversity.

Moreover, the enhanced cell type classification can be substantiated by the gene modules that were identified by HotSpot [11] (see Additional file 4: Table S1 for top module genes). In the uncorrected MPs data, Module 2 (which contains a substantial number of KRT genes) and Module 3 (which contains RPL genes) are the most prominent (Figure 3C). However, following correction, no modules based on KRT genes were identified. In contrast to the uncorrected data, the corrected dataset reveals a broader tumor-associated macrophage (TAM)-related gene module, comprising Modules 2, 3, 8, 9, and 10 (Figure 3D). For example, Module 2 displays high expression of RNASE1, C1QA, and CD163, whereas Module 3 exhibits elevated expression of APOC1, CTSB, and TYROBP. This enhancement allows for a more comprehensive comparison of the immunological profiles in human keloids as explored in the study.

### Performance Comparison of Existing Ambient RNA Correction Tools in Mouse Kidney Samples

To showcase CellClear’s performance among existing ambient RNA correction tools, we conducted evaluations using three scRNA-seq replicates from mouse kidney samples [8]. These samples, known for elevated ambient expression levels in Proximal tubule cells (PTs) and specific ambient genes like Slc34a1, were utilized for benchmarking CellClear against others.

Four tools (CellClear, CellBender, DecontX and SoupX) were performed to show their ability in processing corrected ambient genes, and the cell types identified by them were show in Additional file 2: Figure S1A. Both CellClear and CellBender consistently and effectively reduced the expressions of Slc34a1 in non-PT cells across all three samples (Figure 4A). DecontX showed variable performance, effectively reducing Slc34a1 expressions in two of the samples but performing poorly in the other, whereas SoupX appeared to fail to accurately remove non-specific expressions of Slc34a1 in all replicates.

**Figure 4.**
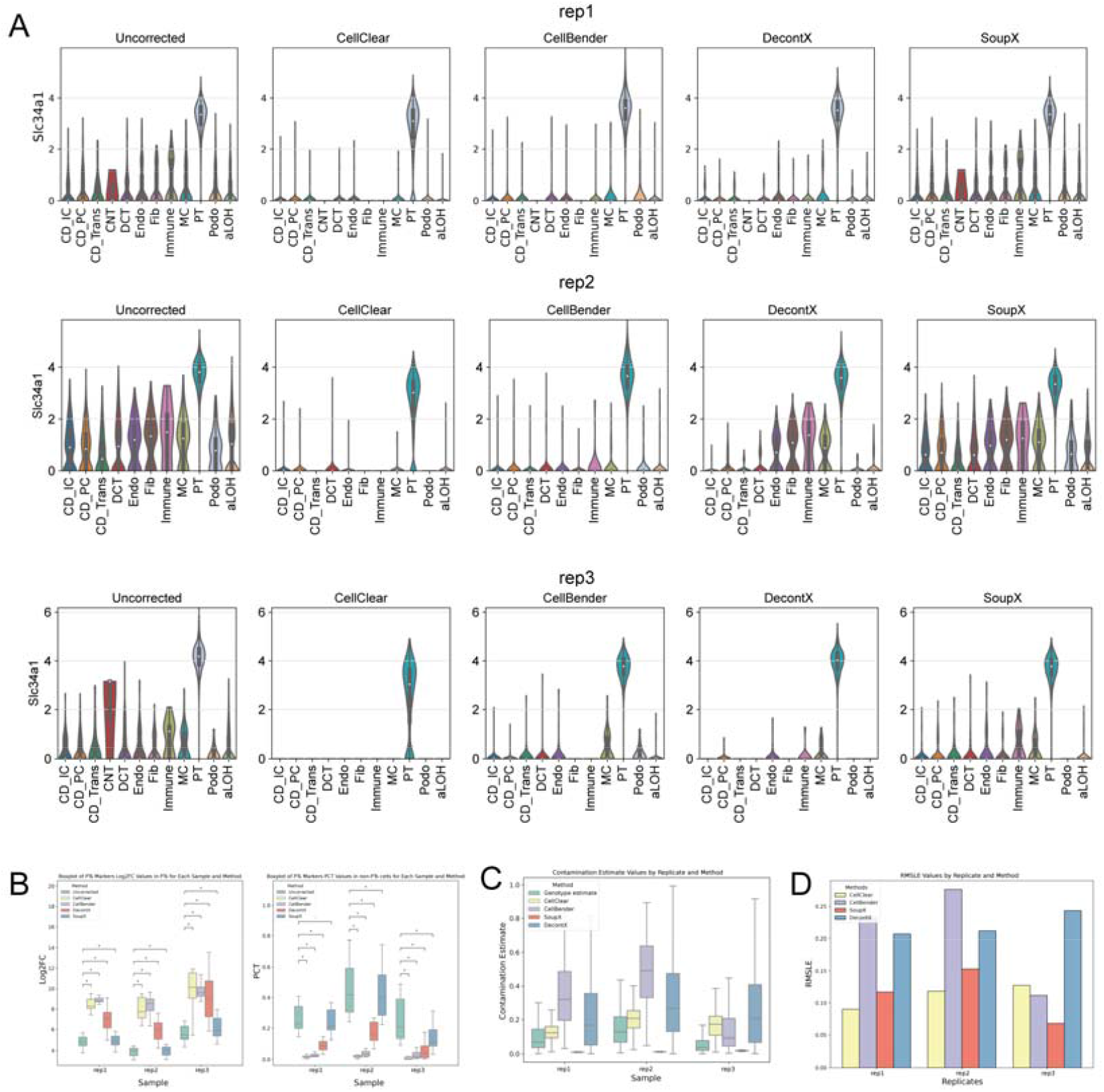
Performance Evaluation of Ambient RNA Correction Tools on Mouse Kidney Samples. (A) Violin plots showing the expression levels of the PTs marker Slc34a1 in three replicates before and after correction by four tools. (PT, proximal tubule cells; CD_IC, intercalated cells of collecting duct; CD_PC, principal cells of collecting duct; CD_Trans, transitional cells of collecting duct; CNT, connecting tubule cells; DCT, distal convoluted tubule cells; Endo, endothelial cells; Fib, fibroblasts; aLOH, ascending loop of Henle cells; dLOH, descending loop of Henle cells; MC, mesangial cells; Podo, podocytes). (B) Comparison of log2FC and percentage of expression (PCT) before and after correction by four tools in differential expression analysis of top 10 PT markers. (C) Estimated ambient expression levels per cell based on genetic variants and four ambient correction tools. (D) Low RMSLE values indicate high similarity between estimated values and the genotype-based estimates.

In the context of ambient gene correction, it is crucial not only to capture ambient genes but also to ensure that the other genes remain unaffected, thereby preserving the biological feature of the data. Moreover, ambient expressions typically are from not just one gene but multiple genes that share similar expression patterns. The ambient expression pattern in this dataset, mainly composed of the top 10 PT markers, was expected to be corrected by four tools, resulting in increased sensitivity of detecting these PT markers. After correction by CellClear, CellBender, and DecontX, the log2 fold changes (log2FC) of the top 10 PTs markers in PT cells versus all other cells were significant increased comparing to the uncorrected data. Meanwhile, both CellClear and CellBender effectively reduced the expression percentage of these markers in non-PTs cells (Figure 4B). For non-ambient genes, the expression levels and percentages of intercalated cells of collecting duct and Endothelial markers remained largely unchanged across all four tools (Additional file 2: Figure S1B).

To evaluate the precision of these tools in detecting ambient RNA expression, the estimated ambient expression fraction per cell provided by each tool were compared, using genotype-based ambient expression estimates in the original publication as the ground truth. As illustrated in Figure 4C, CellClear consistently outperformed the others across the three replicates, highlighting the close alignment of CellClear’s ambient expression fractions in each cell with the genotype-based estimates. Meanwhile, the Root Mean Squared Logarithmic Error (RMSLE) was used to quantify the difference between the estimated background noise fractions per cell from different ambient RNA correction tools and the genotype-based estimates [8], yielding similar conclusions (Figure 4D).

### Advancing Downstream Analysis: A Joint Evaluation of Ambient RNA Correction by CellClear and CellBender

At the end, we sought to understand how corrected data affect downstream analysis, also used by CellBender to demonstrate its performance [5]. In the first human PBMC dataset, the cell type of each cluster in corrected and uncorrected data was identified (Figure 5A) based on the markers provided in the article [5]. CellClear showed an ambient expression level of 0.104 and the top 20 ambient genes included Neutrophil/Monocytes-related genes (S100A8/A9/LYZ), and HLA genes (Additional file 1: Table 1). The widespread expression of Neutrophil/Monocytes-related genes in all cell types of uncorrected data, and their significantly reduced expression in CellClear-corrected data, but to a less degree in CellBender-corrected data (90.5% expressed in non-Monocytes cells versus 0.2% and 7.7%) demonstrate CellClear’s accuracy in identifying and correcting ambient genes (Figure 5B, Additional file 2: Figure S2A, S2B).

**Figure 5.**
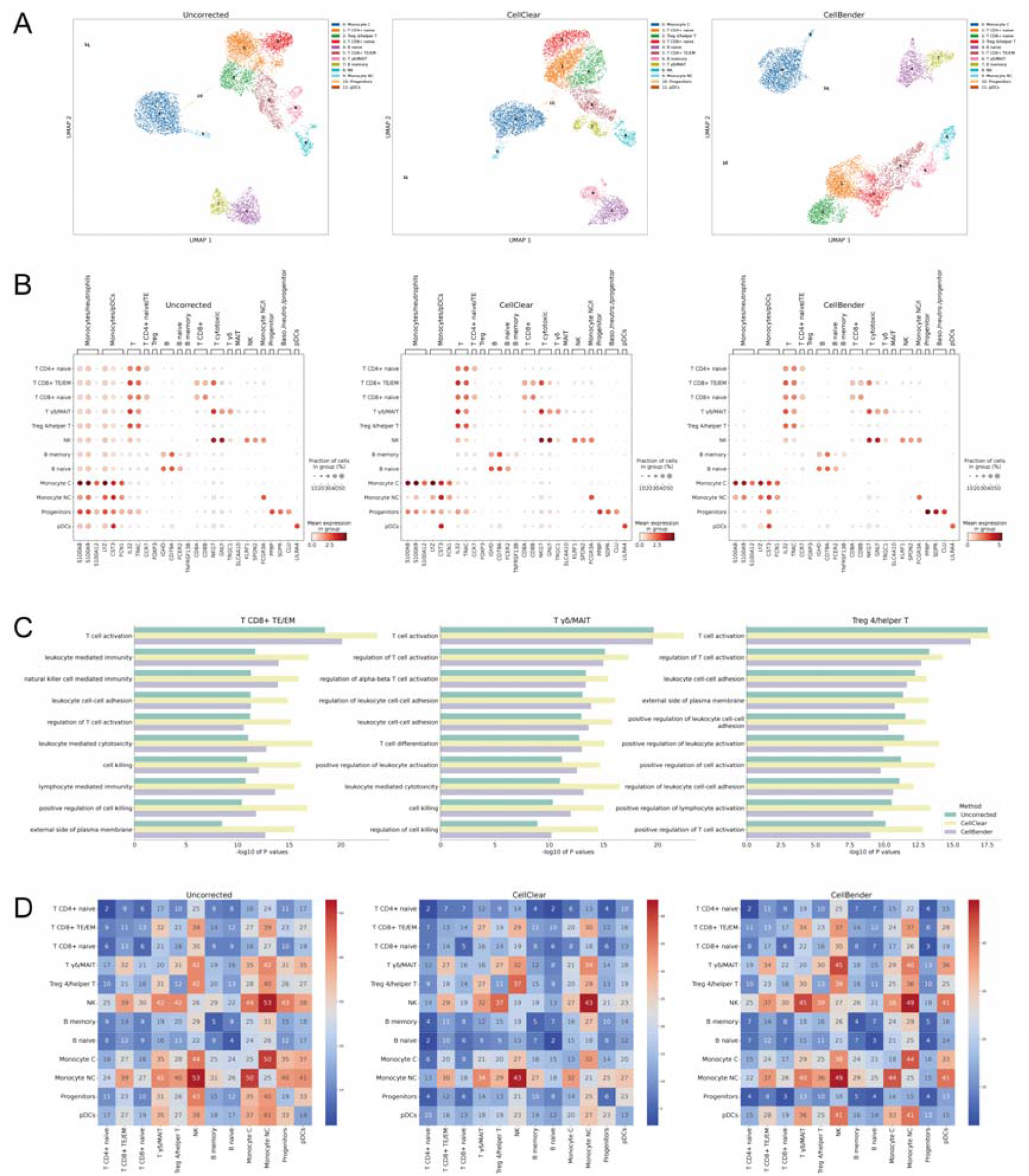
Comparison of the Potential Impacts of Ambient RNA Correction Tools on Downstream Analyses. (A) UMAP scatter plots of a public human PBMC sample before and after ambient expression correction, coloured by cluster labels (Baso., basophil; neutro., neutrophil; NK, natural killer cell; Treg, regulatory T cell; MAIT, mucosal-associated invariant T cell; monocyte C, classical monocyte; monocyte NC, non-classical monocyte). (B) Dot plots showing the expression levels of markers before and after correction. (C) Comparison of -log10 (p-values) for shared top 10 pathway enrichment across each cell type before and after correction, using differentially expressed genes with log2FC ≥ 1. (D) Heatmap showing the number of interaction pairs before and after correction. Color bar from blue to red showed the number of interacting pairs between the two cell types from low to high.

The corrected data from two tools were then used to assess the impact on downstream analyses. Differential gene expression and Gene Ontology enrichment analyses [12] were performed to verify if corrected data could offer a more precise functional interpretation than uncorrected data. Pathway enrichment analyses were compared using the top 10 shared pathways enriched by up-regulated genes from each cell type in uncorrected and corrected data. The analyses showed that in the CellClear-corrected data there was a significant increase in the level of enrichment, particularly for T cell subtypes that were significantly affected by the ambient expression of Neutrophil/Monocyte related genes (Figure 5C). This demonstrates that correction may enhance the sensitivity of pathway analysis for each cell type Cell-cell interactions could also be significantly affected by ambient expression. After correction by CellClear, CellphoneDB [13] analysis revealed a notable decrease in the number of interactions between cell types, with the total number of significant interacting pairs decreasing from 1815 to 1233 (Figure 5D). The data corrected by CellBender showed a similar pattern but to a less degree, decreasing from 1815 to 1564. This result suggested that inflated background noise due to ambient expression could potentially cause some of the interactions to be incorrectly detected by cell communication tools such as CellphoneDB. For example, previous study has shown the interaction pair CD55-ADGRE5 between Monocytes and T cells plays a crucial role in T cell activation [14]. This particular interaction was falsely identified as a pair between B cells and T cells due to Monocytes-related contamination in the uncorrected data, while CellClear corrected data reduced this false positive.

Furthermore, two human testis samples were analyzed using Monocle3 [15] to investigate the impact of contamination correction tools on the developmental trajectory of germ cells. From the dimensionality reduction and pseudotime trajectory plots, it was observed that cells followed the developmental path from spermatogonial stem cells (SSC) to sperm both before and after correction, indicating that ambient RNA expression or its correction does not significantly affect the overall developmental trajectory of the data (Figure 6A,6B). However, Figure 6C shows that after correction with CellClear, the marker genes for each germ cell type exhibited greater stage-specific expression. Additionally, Figure 6D demonstrates that while the overall developmental trajectory remained unaffected, the variance in pseudotime for the same cell type was reduced following CellClear correction, resulting in a reduction in dispersion.

**Figure 6.**
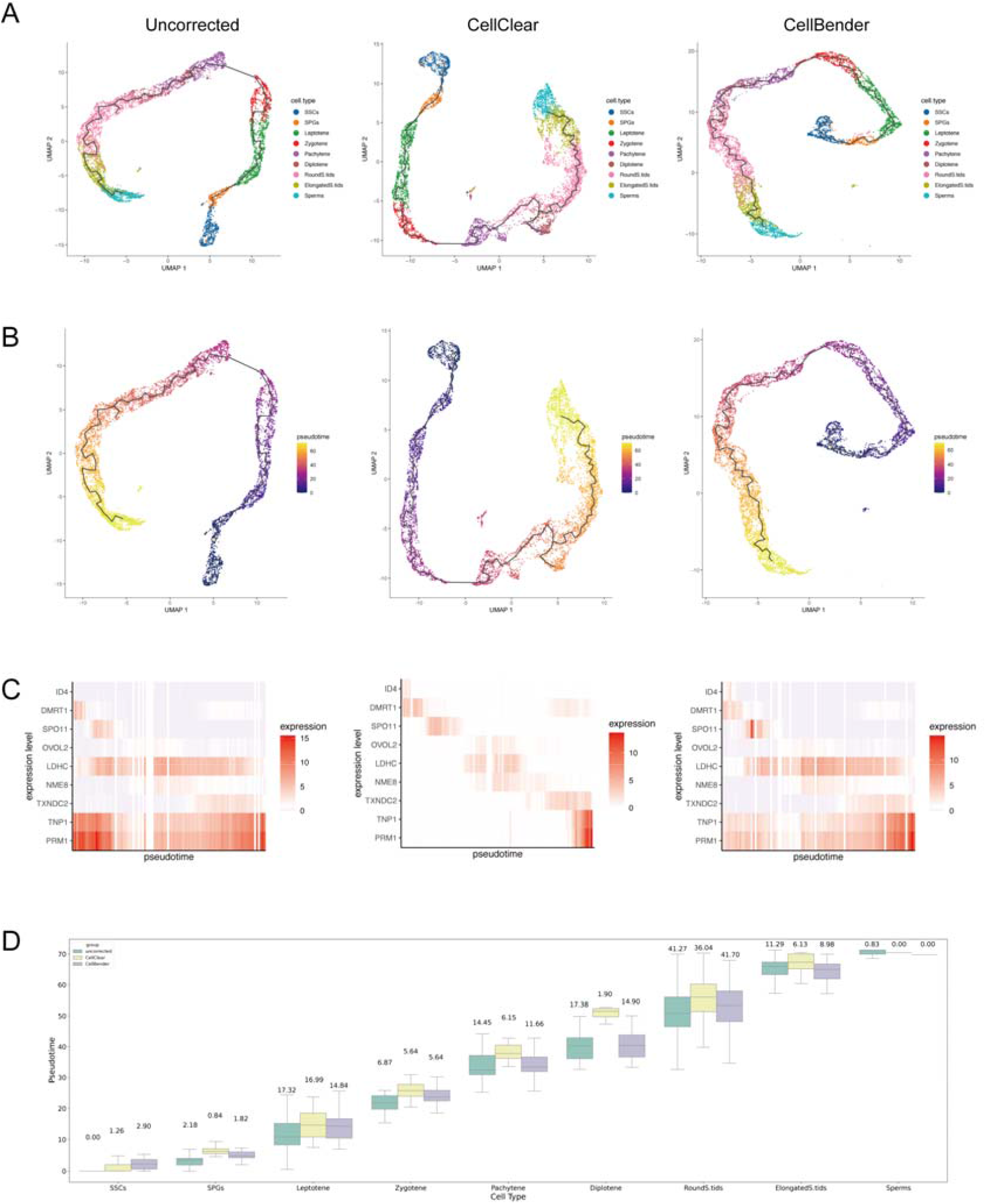
Comparison of Germ Cell Development Before and After Ambient RNA Correction. (A) Dimensionality reduction plots of germ cells before and after correction, coloured by cluster labels (SPGs, spermatogonia; SSCs, spermatogonial stem cells; Leptotene, leptotene spermatocytes; Zygotene, zygotene spermatocytes; Pachytene, pachytene spermatocytes; Diplotene, diplotene spermatocytes; RoundS.tids, round spermatids; ElongatedS.tids, elongated spermatids; Sperms, sperms). (B) Pseudotime trajectory changes of germ cells with and without correction. (C) Expression of marker genes for each germ cell type across pseudotime intervals, before and after correction. (D) Pseudtime distribution for each germ cell type, with variance shown above the boxplots.

In summary, CellClear outperformed other tools in identifying and correcting ambient genes efficiently, preserving the original biological features and improving the accuracy of downstream analysis.

## Conclusions

CellClear distinguishes itself from existing tools like DecontX, SoupX and CellBender by its superior capability to correct ambient genes while preserving biological features. The top ambient genes and their contamination levels provided by CellClear enable researchers to quickly assess the data during the quality control stage, identify the primary contamination sources, and correct them when necessary. Comparing to the discovery of data quality issues related to ambient RNA after the data integration step, this method avoids the tedious work to iterate between QC and data integration. Furthermore, the corrected data demonstrated cell types with more specific differentially expressed genes and significantly enriched key pathways, revealing its capability for enhancing downstream analysis. All these evidences substantiate the assertion that CellClear is a reliable tool for restoring true expression programs by selectively removing ambient noises.

## Methods

### Identify background barcodes that exhibit similar expression programs to the foreground clusters

The CellClear method employs clustering and Non-Negative Matrix Factorization (NMF) to derive cluster-relevant expression programs from foreground cell matrix, which is the cell associated matrix identified by primary analysis pipelines. The W and H matrices of NMF are initially set using the cellular identities of individual cells and the log fold changes of differentially expressed genes identified by Scanpy [16]. A clustering resolution range of 0.8 to 1.2 is recommended to obtain more cluster-relevant expression programs. This approach allows us to seed the model with prior information (cluster-specific programs), reduce the risk of overfitting during training and guide the model towards biologically relevant results [17].

Next, the unit variance scaled foreground counts matrix *V*_*km*_, which has m cells and k features, is infinitely approximated with 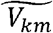:

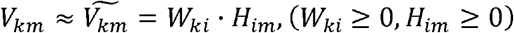

*W*_*ki*_ is the basis matrix, with each row representing a basis vector or an inherent component of the original data. *H*_*im*_ is the coefficient matrix, and each column of matrix *H*_*im*_ indicates the coefficients corresponding to the basis vectors in *W*_*ki*_. The index i represents the number of cluster-relevant expression programs.

The background count matrix is derived from all barcodes captured in the experiment, specifically those with fewer than 100 UMIs, ensuring no cells are included. In a contaminated scRNA-seq experiment, the foreground barcodes primarily contain the native gene expression program A, the contaminated marker’s gene expression program B, and the non-specific ambient gene expression program C (such as housekeeping genes, etc.). Programs B and C together determine the ambient expression in each sample. Program C exhibits a relatively uniform distribution and a minimal impact across most of the data, while the expression of Program B varies in each sample, primarily influenced by the number and diversity of contamination sources. If background barcodes exhibit a similar expression program to one of the foreground clusters, it implies that these background barcodes share a potential program B with their corresponding foreground clusters.

To identify background barcodes with expression programs similar to foreground clusters, two step Non-Negative Least Squares (NNLS) approximation is applied to background counts matrix:

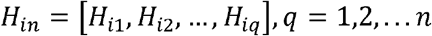

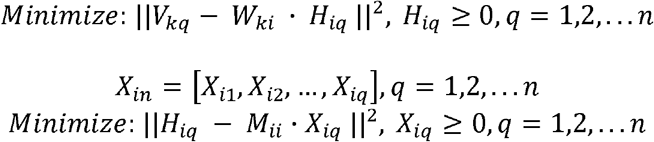

In first step of NNLS, *V*_*kn*_ is the unit variance scaled background counts matrix which has k features and n barcode. For each column of *V*_*kn*_, which can be denoted as *V*_*kq*_, NNLS seeks to find a non-negative coefficient vector *H*_*iq*_ that best approximate *V*_*kq*_ as a linear combination of the basis vectors in *W*_*ki*_. *H*_*iq*_ is calculated independently and finally forms the non-negative coefficient matrix *H*_*in*_, which represents the distribution or the usage of each background barcode with respect to the ith program. In the second NNLS step, foreground cells within the same cluster are grouped together, and the median value of *H*_*im*_ for each program is calculated within each cluster, denoted as *M*_*ii*_ By minimizing the above equation for each column q independently, a non-negative coefficient matrix *X*_*in*_ can be achieved, representing the contribution of each cluster expression program to each background barcode. Only background barcodes with a maximum proportion value larger than 0.95 are considered for subsequent steps, which means their expression programs closely match with one of the foreground clusters.

### Identification of potential seed ambient genes

We next evaluate the ambient expression level and identify seed ambient genes using a smooth spline method with clusters sharing similar expression programs with at least one background barcode. As the ambient expression program has been divided into two types, we observed that cell type specific contamination program B show a discrete gap between background barcodes and their corresponding foreground clusters while non-specific contamination program C doesn’t. The considerable difference between expected and actual gene expression compared to other genes will help to distinguish ambient genes in program B comparing to the others.

For each cluster *c*, we separately calculate average expression values for all genes g in background barcodes *d* and foreground barcodes *r* identified by the above step, denoted as 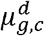 and 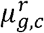 respectively. Each run of fitting involves randomly selecting genes and using the UnivariateSpline function in Python’s scipy package to obtain the predicted average expression level of the genes in the background based on the foreground values. The average gene expressions in the foreground barcodes, are sorted in ascending order to ensure the data is an increasing vector.

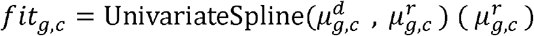

The distance of fitted values to actual average background expression can be denoted as:

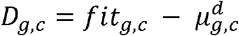

To ensure the stability of the model and the comprehensiveness of the fitting results, we use 60% of the genes for each fitting and perform 100 iterations. Genes with the highest *D*_*g,c*_, are more likely to be potential ambient genes.

We then pooled the gene-*D*_*g,c*_ pairs from each cluster together and the top 20 genes, sorted by *D*_*g,c*_, are considered as potential seed ambient genes. Additionally, the average value of the top 20 genes’ *D*_*g,c*_ is employed as the overall ambient level of the sample. A higher level signifies that the sample is severely affected by ambient expressions.

### Correction of ambient genes

The final step is to capture all potential ambient genes based on the seed ambient genes according to their contribution to gene expression program *W*_*ki*_ and find an ambient expression cutoff to correct them.

Potential ambient expression programs typically are from few source ambient clusters and contaminate other clusters. By examining the median coefficient value of barcodes from the *H*_*im*_ matrix in each cluster, we can determine the weight of each program to the clusters. The program with the highest weight in the greatest number of clusters is considered the potential ambient expression program.

Next, genes with non-zero contribution to this ambient program are identified, and their Pearson’s correlation with the previously identified seed ambient genes is used to capture more ambient genes. A gene is considered as a potential ambient gene if it has a correlation of 0.95 or higher with the previously identified seed ambient genes. The clusters where this ambient expression program has the highest weight are considered potential source ambient clusters.

Binary classification will then be used twice for each ambient gene: (1) identify all potential ambient source clusters (2) determine the ambient gene expression threshold.

Firstly, for each ambient gene, cells in the cluster with the highest average expression are assigned a label of 1. Cells in the remaining clusters are assigned a label of 0. We aim to use AUC-ROC here to determine if some clusters with a label of 0 cannot be distinguished from cells with label 1, thereby identifying additional potential source clusters of this particular ambient gene. If the obtained AUC-ROC value is less than or equal to 0.6, it indicates that the 0 and 1 clusters share a similar gene expression pattern.

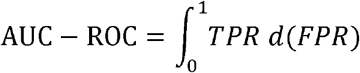

TPR is short for the true positive rate, FPR is short for the false positive rate.

Secondly, we relabel the indistinguishable clusters identified above as 1, and binary classification can be performed again. Youden’s J statistic is used to determine the best threshold of correctionfor each ambient gene.

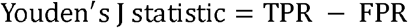

By subtracting this count threshold from the expression values and ensuring non-negative values, the final corrected expression values are obtained.

### Existing Ambient RNA correction tools

For CellBender (v0.3.0), remove-background module was used to run on the raw matrix, with the default recommended settings (fpr 0.01, epochs 150). The parameter expected-cells and total-droplets-included was set to 3,000 and 15,000.

For DecontX (v1.16.1) and SoupX (v1.6.2), raw matrix and filtered matrix were used as input. Cluster labels for filtered matrix were generated by Louvain (0.8.1) on 30 principal components and a resolution of 1 as implemented by FindClusters in Seurat (v4.3.0.1) after normalization and feature selection of 3000 genes. The ambient RNA correction processes for DecontX and SoupX both utilized default parameters.

## Data availability

The public datasets study in this article include human-mouse cell mixture data, kidney scRNA-seq data, PBMC scRNA-seq data, and human skin scRNA-seq data. The contamination metrics and top contamination genes for the remaining samples are provided in the Additional fil e 1: Table S1.

The human-mouse cell mixture data is available at https://www.10xgenomics.com/datasets/12-k-1-1-mixture-of-fresh-frozen-human-hek-293-t-and-mouse-nih-3-t-3-cells-2-standard-2-1-0. For this sample, we removed the top 5% of barcodes with the highest UMI counts and unique gene counts, and excluded barcodes with a mitochondrial read fraction beyond 10%.

The kidney scRNA-seq data, with annotation and genotype information, is available at https://zenodo.org/records/7328632. For the three kidney samples, we removed the top 2% of barco des with the highest UMI counts and unique gene counts, and excluded barcodes with a mitochondrial read fraction beyond 50% and gene counts below 200.

The PBMC scRNA-seq data is available at https://www.10xgenomics.com/resources/datasets/8-k-pbm-cs-from-a-healthy-donor-2-standard-2-1-0. For this sample, we applied the same fil tering criteria as for the human-mouse cell mixture data.

The human skin scRNA-seq datasets were obtained from NCBI Sequence Read Archive at P RJNA844167 and processed using the same criteria as the kidney scRNA-seq data.

The human testis scRNA-seq datasets were obtained from NCBI Sequence Read Archive at P RJNA615042 and processed using the same criteria as the kidney scRNA-seq data.

Dimensionality reduction and clustering analysis for all datasets were performed using Scanpy (v1.9.3).

## Processing Temporal and Spatial Variations in 35 Single-Cell Samples

In terms of resource consumption, both runtime and memory usage increase significantly as the data size grows, exhibiting a nearly linear relationship. (Additional file 2: Figure S3A).

## Code availability

CellClear and the code used in this study is available at https://github.com/WhiteRabBio/CellClear under the BSD 3-Clause License.

## Key Points

- CellClear estimates the overall ambient level and identifies potential ambient genes for each sample efficiently, enabling researchers to assess contamination related data quality during scRNA data preprocessing.
- CellClear locates ambient-specific expression programs to correct, minimizing the unnecessary distortion of the native information of individual cells. Two example datasets were used to showcase it performs better than state-of-art tools currently available.
- CellClear-corrected data can improve the accuracy of downstream analysis such as pathway analysis and cellular interaction analysis.

## Supplementary Information

## Additional file 1

Table S1. Summary of analyzed scRNA-seq and snRNA-seq datasets.

## Additional file 2

Figure. S1. Performance evaluation of existing tools on mouse kidney samples.

Figure. S2. Performance evaluation of CellClear and CellBender on a human PBMC sample.

Figure. S3. Relationship between Resource Consumption and Data Size.

## Additional file 3

Table S1. Metrics obtained after applying CellClear to correct the “species-mixing” dataset, including the top-expressed mouse genes detected in human cells and the top-expressed human genes detected in mouse cells.

## Additional file 4

Table S1. Significant genes associated with each MP module before and after correction.

## Declarations

### Ethics approval and consent to participate

Not applicable

### Consent for publication

Not applicable

## Acknowledgements

Not applicable

