## Additional file 2 for "CellClear: Enhancing Single-cell RNA Data Quality via Biologically-Informed Ambient RNA Correction"

**Additional file 1**

**Supplementary Figure 1**

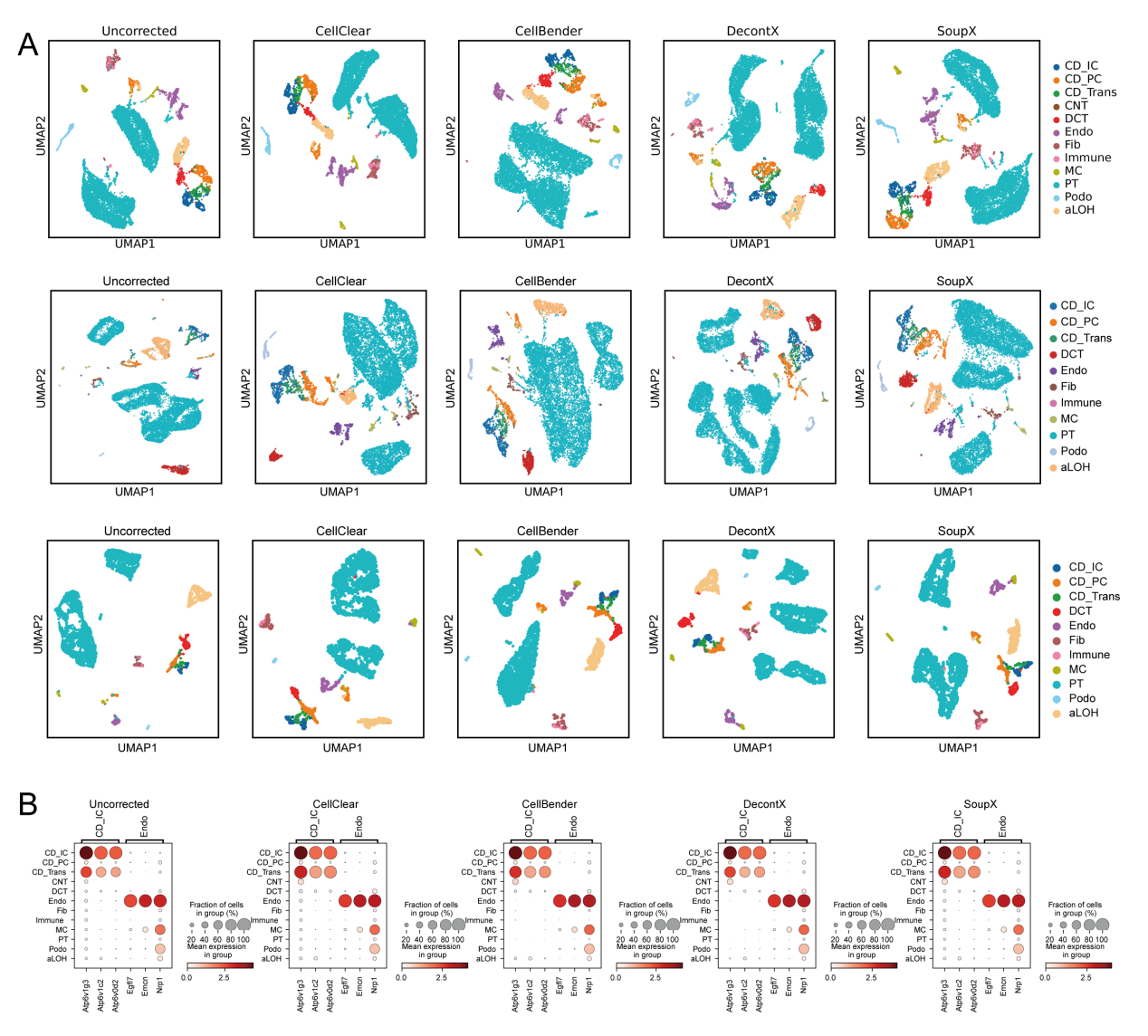

**Figure. S1 Performance evaluation of existing tools on mouse kidney samples.**

(A) UMAP scatter plot of mouse kidney replicates before and after ambient expression correction, colored by cell type labels. (B) Dot plot depicting the expression levels and percentages of CD_IC and Endo markers in replicate 1.

**Supplementary Figure 2**

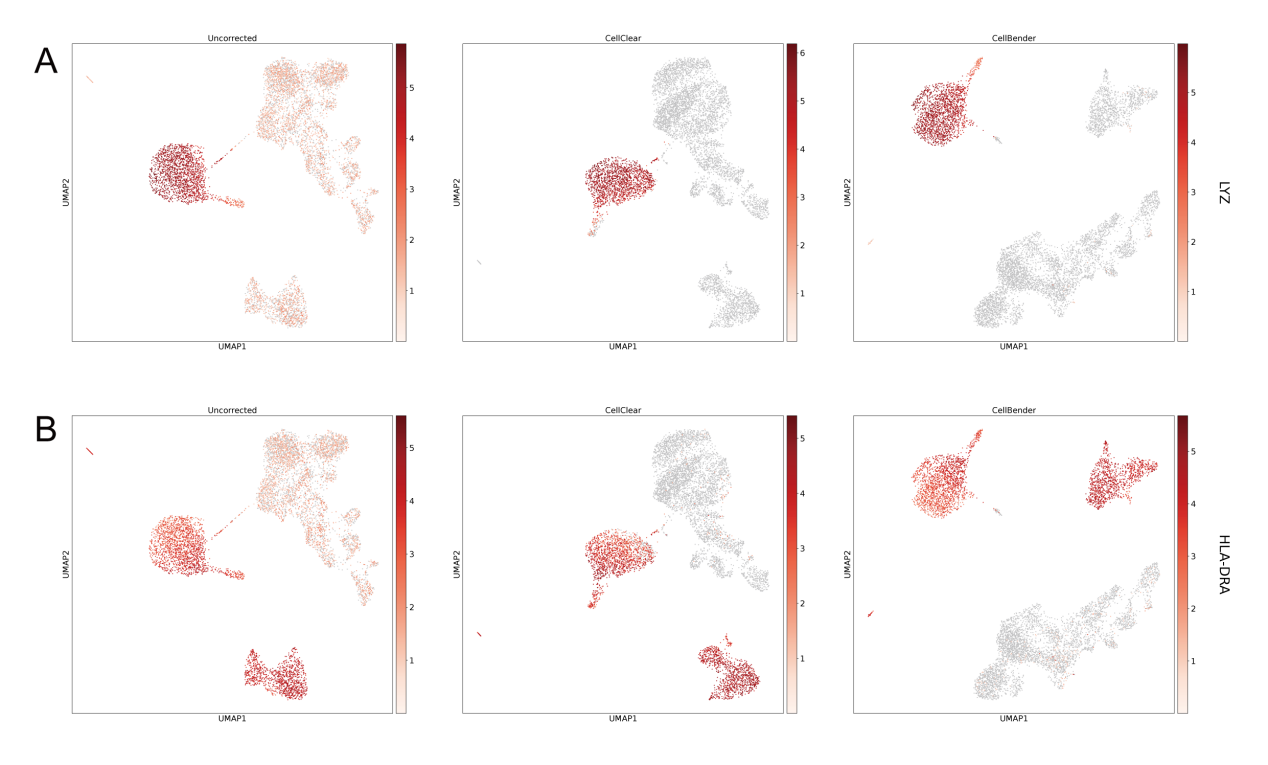

**Figure. S2 Performance evaluation of CellClear and CellBender on a human PBMC sample.**

(A, B) UMAP scatter plots of the expression of LYZ and HLA-DRA in each cell before and after correction.

**Supplementary Figure 3**

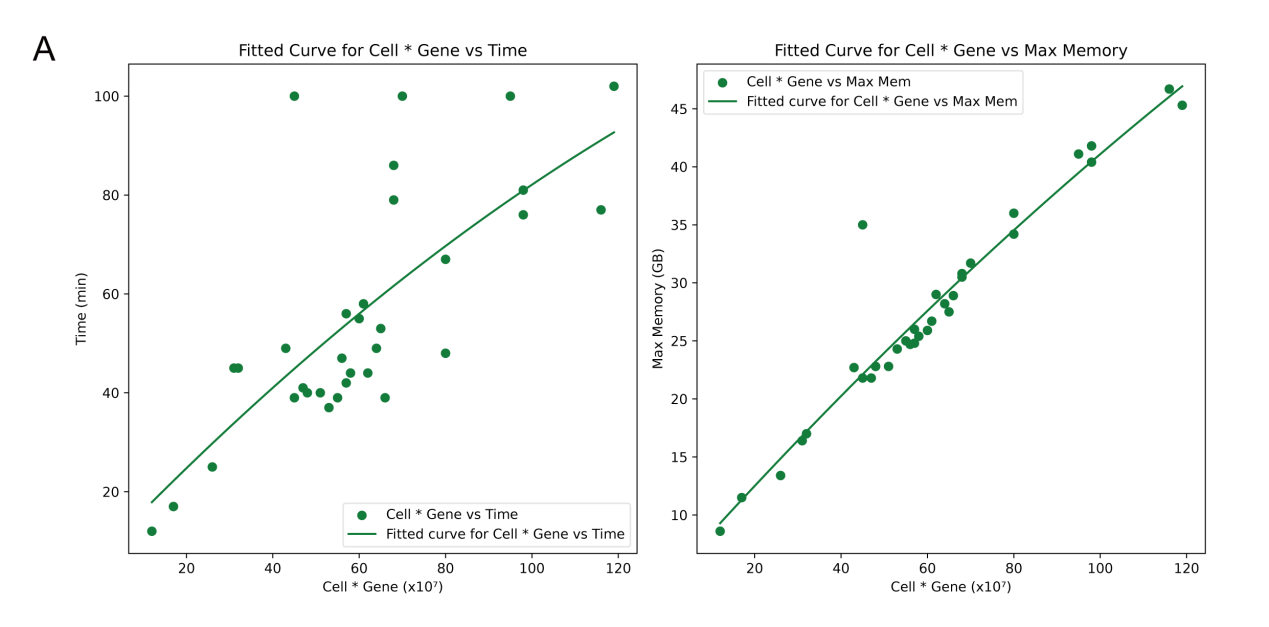

**Figure S3. Relationship between Resource Consumption and Data Size**

(A) The left plot shows the relationship between the cell-gene product and computation time. Green dots indicate data points representing the cell-gene product versus time, and the fitted curve reveals the trend. The right plot illustrates the relationship between maximum memory usage and the cell-gene product. Green dots represent the cell-gene product versus maximum memory usage, with the fitted curve highlighting the trend.
